# Few-Shot Classification of *C. elegans* Developmental Stages via Explainable Hierarchical Hyperbolic Graph Embeddings

**DOI:** 10.64898/2026.06.21.733631

**Authors:** Nazish Khalid, Logan Elliott, Tayo Obafemi-Ajayi, Donald Wunsch, Andrea Scharf

**Affiliations:** Department of Electrical & Computer Engineering, Missouri University of Science and Technology, MO, USA; Department of Biological Sciences, Missouri University of Science and Technology, MO, USA; Engineering Program, Missouri State University, Springfield, USA

**Keywords:** Hyperbolic learning, few-shot classification, developmental staging, *Caenorhabditis elegans*, interpretability, explainability

## Abstract

Automated, accurate, and fast developmental-stage classification of *C. elegans* from microscopy-based morphological images is essential for aging research, drug screening, and disease modeling. However, it remains challenging due to morphological similarities between stages and the limited annotated data. In this work, we propose HyperDev, a hyperbolic few-shot learning framework that addresses these limitations by directly encoding developmental hierarchies in the embedding space, unlike conventional Euclidean approaches that treat stages as independent classes. HyperDev uses Poincaré ball geometry, combined with a biologically informed developmental prior, to naturally represent stage relationships. We introduce our self-curated *C. elegans* dataset spanning seven developmental stages (Egg, L1–L4, Adult, Dauer) with extreme class imbalance (6-8 samples per minority class). HyperDev achieves competitive classification accuracy (76.9–88.3%) while providing intrinsic explainability across nine 7-way few-shot evaluation settings. The learned embeddings exhibited strong biological alignment (Pearson *r* = 0.669, *p <* 0.001), while significantly outperforming ProtoNet (*r* = 0.187), MatchingNet (*r* = 0.235), and RelationNet (*r* = 0.464). These results establish hyperbolic geometry as a principled approach to explainable few-shot learning in biological imaging, where understanding learned representations is as critical as predictive performance.

**Clinical Relevance:** By enabling explainable, data-efficient developmental staging from scarce samples, HyperDev supports improved phenotype quantification for aging research, disease modeling, and drug screening.

## I. Introduction

Due to severe class imbalance, high morphological similarity between adjacent stages, and the need for explainable models that align with the biological knowledge, the development of different stage classification from microscopic images remains a significant challenge. The model organism *Caenorhabditis elegans* (*C. elegans*) progresses through seven developmental stages/states: egg, L1–L4, adult, and dauer [1], automated image-based stage classification remains challenging due to the reliance on expert-driven manual annotation and the time-intensive nature of generating high-quality labeled data. The difficulty of this task lies in its time-consuming nature (30-60 seconds per sample), requires substantial morphological expertise, and does not scale to modern high-throughput imaging platforms capable of producing hundreds of thousands of image per experiment. Moreover, existing classification methods ignore the inherently hierarchical structure of development, treating stages as independent classes despite their biological lineage relationships. Accurate identification of developmental stages is a cornerstone of *C. elegans* research and is essential for a wide range of studies, including aging [2], neurodegenerative disease modeling [3], and high-throughput drug screening [4]. The importance of this organism stems from its exceptionally well-characterized biology, including its fully mapped 959-cell lineage [5], highly conserved genetic pathways [6], and its role in foundational discoveries such as apoptosis [7] and RNA interference [8]. Reliable stage classification allows researchers to align observed phenotypes with specific developmental time points, identify disruptions in developmental timing in disease models, and study stage-dependent drug responses. However, reliance on expert manual annotation limits throughput and introduces subjectivity, particularly near stage boundaries, while existing automated methods lack interpretability and fail to capture biologically meaningful developmental relationships. A principled, explainable framework that maintains accuracy under data scarcity is therefore critical for advancing developmental biology research.

Existing few-shot learning methods perform well in low-data settings but struggle with hierarchical biological problems. Euclidean metric-based and meta-learning approaches [9] treat classes symmetrically and lack inductive bias to capture developmental hierarchies, where adjacent stages should be more similar than distant ones. While hyperbolic neural networks naturally model hierarchical structure [10], [11], they have not been explored in few-shot biological settings requiring interpretability and robustness to class imbalance. As a result, no existing approach jointly addresses few-shot learning, hierarchical representation, and biological interpretability for developmental stage classification. Recent work in computer vision has further demonstrated that Euclidean and spherical embedding spaces are suboptimal for hierarchical representation learning, and that hyperbolic geometry naturally captures such structure in open-world settings [12].

We present *HyperDev*, a hyperbolic few-shot learning framework that addresses these limitations by exploiting the natural alignment between hyperbolic geometry and hierarchical biological structure. Developmental stages are modeled as nodes in a phenotypic progression hierarchy, embedded in the Poincaré ball using a ResNet-18 visual encoder [13] with exponential mapping, where geometric distances encode stage similarity and radial position reflects developmental commitment, with larger radius corresponding to increased specialization. To incorporate domain knowledge, HyperDev introduces a developmental prior attention mechanism that weights class prototypes according to known biological proximity, providing a principled inductive bias beyond generic attention [14]. The presented technique yields explainable embeddings with biological meaningful geometry that enables the visualization of the developmental structure and shows us the improved understanding of model decision under data collection limitation.

### Contributions

The main contribution of this work is as follows:

- **HyperDev Framework:** We propose HyperDev, a few-shot learning-based developmental stage classifier that combines Poincaré ball embeddings with a biologically informed developmental prior attention mechanism.
- **Benchmark Dataset:** We created a challenging *C. elegans* open-source microscopy dataset with severe class imbalance (6–8 samples per minority class), enabling evaluation of few-shot learning methods in a realistic biological setting. The dataset used in this study is publicly available through Missouri S&T Scholars’ Mine at https://scholarsmine.mst.edu/research_data/16/.
- **Explainability and Performance:** The classification results of HyperDev show 76.9–88.3% accuracy, but the method produces better biological alignment than baseline methods (*r* = 0.187–0.464) because it achieves a Pearson correlation of (Pearson *r* = 0.669, *p <* 0.001). The learned embeddings show developmental stages in a sequence that matches biological development patterns, and most prediction errors occur between adjacent stages.
- **Evaluation Protocols:** The research provides comprehensive evaluation methods for few-shot developmental biology, spanning assessment protocols for hierarchical correlation, geometric structure analysis, and biologically grounded error characterization.

Our results demonstrate that hyperbolic geometry provides a principled and explainable representation space for few-shot learning in developmental biology, where understanding model behavior is as critical as predictive performance. Fig 1 explains the pipeline of whole framework.

**Fig. 1:**
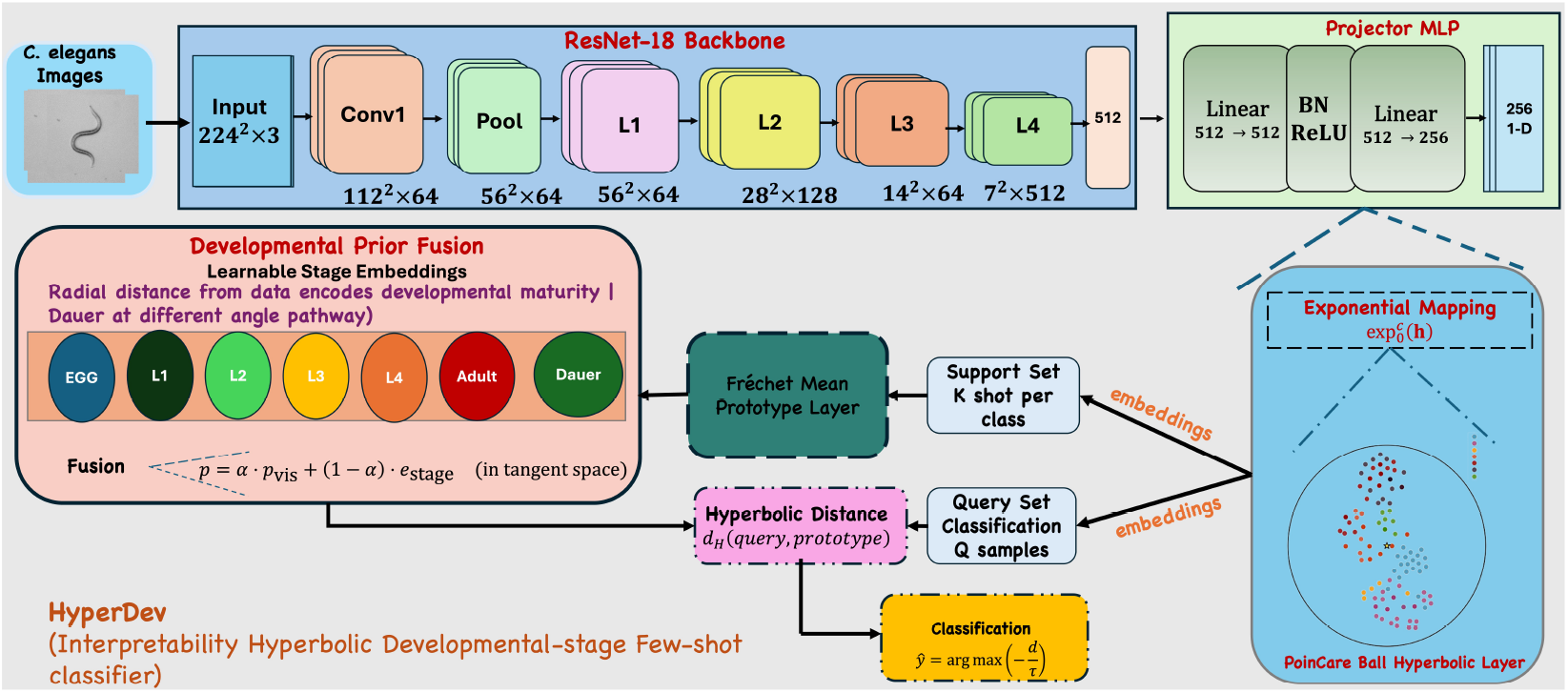
The Poincaré ball contains embedded images that use hyperbolic geometry to represent the hierarchical development pattern of *C. elegans*. Class prototypes are computed using the Fréchet mean, presenting stable, geometry-aware few-shot representations, and queries are classified using hyperbolic distances to these prototypes. The curvature of the Poincaré ball separates early and late stages by expanding representational capacity toward the boundary, preserving fine-grained developmental transitions that Euclidean spaces struggle to model. The geodesic aggregation method reduces model sensitivity to noise when it works with limited supervised training data. The prototype-based attribution method produces explanations that people can understand and that stem from the system’s actual structure. The system uses these elements to create a method that classifies developmental stages through few-shot learning while maintaining biological relevance.

## II. Methodology

### A. Data Collection and Annotation

We contribute a *C. elegans* developmental stage dataset comprising 411 microscopy images with 651 annotated instances across seven stages (Egg, L1–L4, Adult, Dauer). Images were acquired using a Nikon SMZ18 stereomicroscope (10× magnification) from wild-type N2 liquid cultures sampled at strategic time points: Day 1 (L1), Day 2 (L2), Day 3 (L2/L3), Day 4 (L4/Adult), and Day 5 (Adult) [15], [16]. Dauer larvae [17] were obtained via sodium dodecyl sulfate resistance assays. Instance segmentation annotations were created with assistance from the Segment Anything Model 2 and stored in COCO format using open source digital sreeni software [18]. Stage assignments follow established morphological criteria and developmental timing from WormAtlas [1], [19]: newly hatched (L1), intestinal visibility (L2/L3), vulval slit (L4), and embryo presence (Adult). The L1/L2 and L2/L3 boundaries present inherent ambiguity due to continuous development without discrete anatomical transitions. Fig 2 shows the hardware setup for dataset collection.

**Fig. 2:**
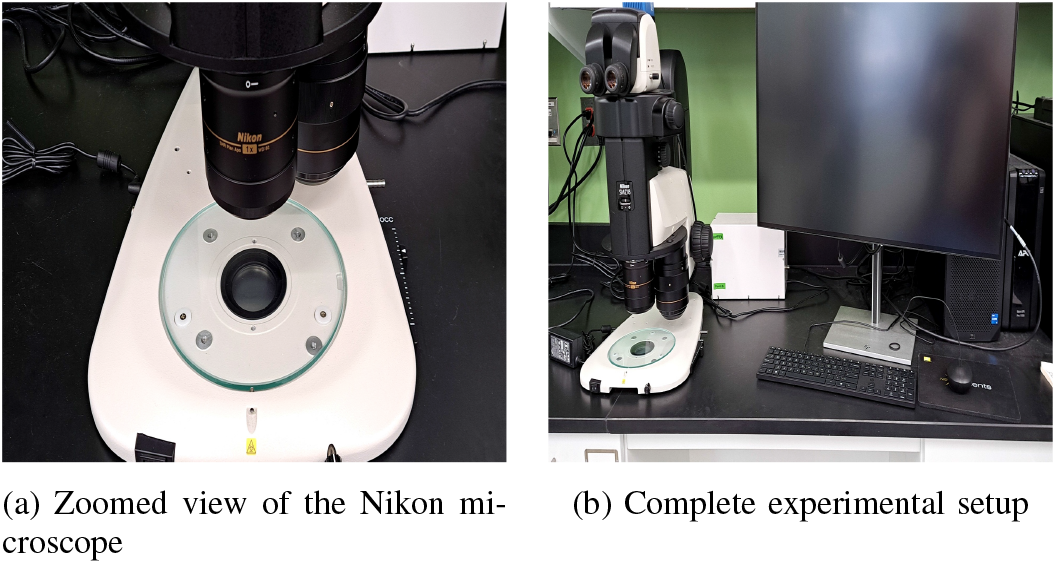
Hardware setup used for dataset acquisition using the Nikon SMZ18 microscope at 10*×* magnification.

### B. Dataset Characteristics

The collected dataset is highly imbalanced, with a large number of early larval samples (Egg: 131, L1: 148, L3: 128), fewer instances from later reproductive stages (L4: 93, Adult: 82), and very limited examples of transition stages (L2: 30, Dauer: 39). In particular, the L2 stage is represented by only 30 training images, which reflects its short developmental duration (approximately 8–12 hours at 20 ^°^C) and its strong morphological resemblance to neighboring stages [20]. This realistic scarcity makes L2 and Dauer especially challenging cases and well suited for evaluating few-shot learning approaches. *C. elegans* grown in liquid medium are thinner and longer than on solid media, which complicates stage identification.

#### Dataset Preprocessing

Images were resized to 256×256 pixels and center-cropped to 224×224 to match standard ImageNet-pretrained architectures. Instance crops were extracted using COCO bounding boxes with 10% padding. All images were normalized using ImageNet statistics. During training, we applied standard augmentations (horizontal/vertical flips, ±15° rotation, color jitter) with additional elastic deformations to simulate natural body contortions. Augmentation intensity was increased for underrepresented classes (L2, Dauer) to address overfitting.

### C. Model Architecture

We address seven-way developmental stage classification of individual *C. elegans* instances (Egg, L1–L4, Adult, Dauer) under a few-shot regime. HyperDev follows an episodic prototypical learning paradigm and embeds visual representations in a hyperbolic space structured to reflect the hierarchical nature of worm development.

#### 1) Problem Formulation

Few-shot learning is performed using *N*-way, *K*-shot episodes. Each episode samples a support set presented by equation 1

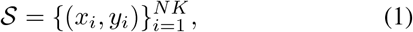

and a query set by equation 2

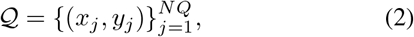

where *y* ∈ *Y* = {1, …, 7}. We use *N* = 7, *K* = 5, and *Q* = 15 throughout. Evaluation episodes are drawn from held-out individuals to assess generalization.

#### 2) Backbone and Feature Projection

Each image *x* is encoded using a ResNet-18 backbone pretrained on ImageNet with the classification head removed. Global average pooling yields a 512-D feature vector, which is projected to a *d* = 256 dimensional embedding via a two-layer MLP with batch normalization and ReLU activation presented by equation 3:

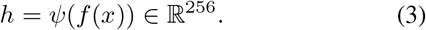

#### 3) Hyperbolic Embedding

Projected features are mapped into a *d*-dimensional Poincaré ball 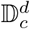 with curvature *c* = 1 using the exponential map at the origin. This mapping allocates increasing representational capacity toward the boundary, which is well suited for hierarchical developmental structure. Hyperbolic distances between embeddings are computed using the Poincaré metric.

#### 4) Hyperbolic Prototypical Classification

For each class *k*, support embeddings are mapped to the tangent space, averaged, and projected back to hyperbolic space to form a class prototype shown by equation 4:

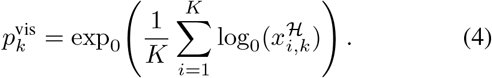

Query samples are classified based on hyperbolic distance to prototypes, scaled by a learnable temperature and optimized using cross-entropy loss.

#### 5) Developmental Prior and Hierarchy Regularization

A central design choice in HyperDev is the explicit integration of biological prior knowledge about the developmental trajectory into the hyperbolic embedding space. We introduce learnable stage-specific embeddings presented by equation 5

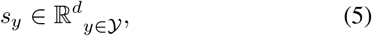

initialized to reflect known developmental structure, with decreasing radial distance from Egg to Adult and a separate off-path branch for Dauer. After projection, these form developmental prior embeddings 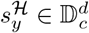.

For each class *k*, the visual prototype 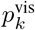 is fused with its corresponding developmental prior 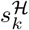 via a learnable interpolation in the tangent space:

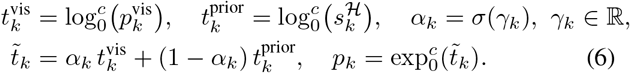

where *γ*_*k*_ is a learnable, stage-specific parameter. This formulation allows prototypes to adaptively balance visual evidence and biological priors.

To enforce biological consistency, we regularize the priors to follow the known developmental ordering (Egg → L1 → L2 → L3 → L4 → Adult). Let 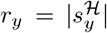 denote the radial norm of stage *y*. We impose a hinge-style constraint presented equation 7

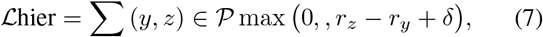

with margin *δ* = 0.08.

This regularization utilizing the exponential geometry of hyperbolic space, where radial position encodes developmental commitment: early stages lie near the boundary, mature stages concentrate toward the origin, and Dauer occupies a distinct high-radius branch, yielding an explainableand biologically grounded embedding structure.

#### 6) Training Objective

The final training objective combines episodic classification loss with hierarchy regularization presented by equation 8:

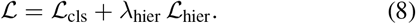

Models are trained episodically using AdamW with gradient clipping and early stopping. Ablation variants are obtained by disabling hyperbolic embedding or hierarchy components while keeping the training protocol fixed.

## III. Experimental Setup

### A. Few-Shot Evaluation Protocol

We evaluate our method under a 7-way, K-shot episodic classification setting that reflects the practical constraints of developmental biology, where only a handful of labeled images are available per stage. Each episode consists of *N* = 7 classes, *K* labeled support instances per class, and *Q* = 15 query samples per class. Episodes are sampled independently for training, and testing using a balanced episodic sampler that ensures class-uniform sampling despite the underlying dataset imbalance. Table I reports dataset statistics for the stratified training, validation, and test splits (60%/20%/20%). During training, we generate 1,500 episodes with random class subsets and instance draws. At test time, we report average accuracy over 500 independently sampled episodes using the exact same 7-way K-shot protocol. All evaluation episodes are constructed from *held-out worms* to prevent identity leakage between splits. However, two minority classes impose a hard constraint on episodic sampling: L2 (6 test instances) and Dauer (8 test instances). For configurations where K+Q>6, strictly disjoint support and query sets cannot be guaranteed for these classes, necessitating sampling with replacement. Specifically, five of nine evaluated configurations are affected (all 15-query settings, 2-shot/5-query, and 5-shot/5-query); results for these are reported separately in Table III (Panel B) and should be interpreted with caution. The four fully held-out configurations (Panel A) serve as the primary basis for comparison.

**TABLE I:** Statistics of the Curated Dataset. The training, validation and test splits are presented while highlighting the imbalance in minority classes.

| Split | Imgs | Inst. | Egg | L1 | L2 | L3 | L4 | Adult | Dauer |
| --- | --- | --- | --- | --- | --- | --- | --- | --- | --- |
| Train | 244 | 389 | 79 | 85 | **19** | 81 | 54 | 49 | **22** |
| Val | 79 | 129 | 24 | 30 | **5** | 27 | 19 | 15 | **9** |
| Test | 88 | 133 | 28 | 33 | **6** | 20 | 20 | 18 | **8** |
| **Total** | **411** | **651** | **131** | **148** | **30** | **128** | **93** | **82** | **39** |

**TABLE II:** Ablation study of HyperDev. Validation accuracy (%) on 7-way 5-shot episodes. ✓= enabled, ✗= disabled.

| Configuration | Hyp. | Dev. | Hier. | Dim | $c$ | Acc. | Ep. |
| --- | --- | --- | --- | --- | --- | --- | --- |
| **(A) Geometry & Components** |  |  |  |  |  |  |  |
| Minimal (Euclidean) | × | × | × | 128 | – | 93.07 | 1300 |
| Euclidean + prior | × | ✓ | ✓ | 128 | – | 90.72 | 1100 |
| No dev. prior | ✓ | × | ✓ | 128 | 1.0 | 90.95 | 1300 |
| No hierarchy loss | ✓ | ✓ | × | 128 | 1.0 | 91.20 | 500 |
| **(B) Embedding Dimension** |  |  |  |  |  |  |  |
| dim = 64 | ✓ | ✓ | ✓ | 64 | 1.0 | 91.62 | 900 |
| dim = 128 | ✓ | ✓ | ✓ | 128 | 1.0 | 92.55 | 1100 |
| dim = 256 | ✓ | ✓ | ✓ | 256 | 1.0 | **93.75** | 700 |
| dim = 512 | ✓ | ✓ | ✓ | 512 | 1.0 | 90.65 | 800 |
| **(C) Curvature** |  |  |  |  |  |  |  |
| $c = 0.5$ | ✓ | ✓ | ✓ | 128 | 0.5 | 90.65 | 400 |
| $c = 1.0$ | ✓ | ✓ | ✓ | 128 | 1.0 | 92.55 | 1100 |
| $c = 2.0$ | ✓ | ✓ | ✓ | 128 | 2.0 | 14.20 | – |
| **(D) Backbone** |  |  |  |  |  |  |  |
| ResNet-18 | ✓ | ✓ | ✓ | 128 | 1.0 | 92.55 | 1100 |
| ResNet-34 | ✓ | ✓ | ✓ | 128 | 1.0 | 92.00 | 400 |
| ResNet-50 | ✓ | ✓ | ✓ | 128 | 1.0 | 90.38 | 800 |
| ResNet-101 | ✓ | ✓ | ✓ | 128 | 1.0 | 91.88 | 800 |
| **HyperDev (Ours)** | ✓ | ✓ | ✓ | 256 | 1.0 | **93.75** | 700 |

Configuration specifies which components are active; Hyp. indicates the use of hyperbolic embeddings, Dev. the developmental-stage prior, and Hier. the hierarchy loss. Dim denotes the embedding dimensionality, $c$ the curvature of the Poincaré ball, Acc. the mean validation accuracy, and Ep. the number of validation episodes used for evaluation.

**TABLE III:** Test accuracy (%) on 7-way *C. elegans* classification (500 episodes). Panel A reports fully held-out configurations where support and query sets are strictly disjoint for all classes. Panel B reports configurations requiring sampling with replacement for minority classes (L2: 6 test instances, Dauer: 8 test instances) because *K* + *Q* exceeds available samples; these results should be interpreted with caution and are included for completeness only.

| Method | Backbone | Panel A: Clean (no replacement) |  |  |  | Panel B: Requires replacement |  |  |
| --- | --- | --- | --- | --- | --- | --- | --- | --- |
|  |  | 1-shot/1-q | 1-shot/5-q | 2-shot/1-q | 5-shot/1-q | 2-shot/5-q | 1-shot/15-q | 5-shot/5-q |
| ProtoNet | ResNet-18 | 80.9±14.5 | 82.9±7.5 | 86.6±12.1 | 88.1±11.4 | 87.3±5.7 | 82.2±8.0 | 89.2±5.0 |
| MatchingNet | ResNet-18 | 81.2±14.5 | 82.1±8.0 | 85.3±12.3 | 86.7±11.5 | 86.3±6.2 | 81.6±8.8 | 89.9±5.1 |
| RelationNet | ResNet-18 | 14.3±0.0 | 14.3±0.0 | 14.3±0.0 | 14.3±0.0 | 14.3±0.0 | 14.3±0.0 | 14.3±0.0 |
| **HyperDev** | ResNet-18 | 76.9±14.0 | 78.4±8.0 | 84.2±12.4 | 87.7±11.3 | 85.2±6.3 | 79.7±6.7 | 88.3±5.1 |

### B. Baselines

To contextualize the contribution of hyperbolic geometry and the developmental prior, we compare HyperDev against strong few-shot baselines:

- **Prototypical Networks (ProtoNet)** [21]: Computes Euclidean class prototypes and classifies queries by distance, serving as the canonical metric-based baseline.
- **Matching Networks (MatchingNet)** [9]: Uses attention over the support set with cosine-similarity prediction, evaluating whether contextual attention alone resolves fine-grained stage differences.
- **Relation Networks (RelationNet)** [22]: Learns a parametric distance function via a convolutional relation module, testing whether expressive similarity functions improve discrimination under severe class imbalance.

All baselines use ResNet-18 backbone, identical augmentations, and the same training schedule for fair comparison. We evaluate each method over 500 test episodes, reporting mean accuracy *±* standard deviation.

### C. Evaluation Metrics

Performance is measured as the mean accuracy across episodic test tasks presented by equation 9:

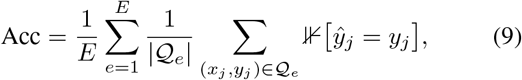

where *E* is the total number of test episodes. We report mean *±* standard deviation over 500 episodes.

To analyze class-specific behavior, we additionally compute per-stage accuracy and a confusion matrix. For interpretability evaluation, we measure the *radial developmental consistency score*, defined as the fraction of stage-prior pairs whose hyperbolic radii follow the expected biological ordering (Egg → Adult), and separately quantify the radial separation of the Dauer branch.

### D. Ablation Studies

Table II systematically isolates the contribution of each HyperDev component. We ablate along four axes: (A) *Geometry & Components* evaluates hyperbolic versus Euclidean prototypes and the impact of the developmental prior and hierarchy loss; (B) *Embedding Dimension* tests capacity scaling from *d* = 64 to *d* = 512; (C) *Curvature* varies the Poincaré ball curvature *c* controlling how sharply distances grow near the boundary; and (D) *Backbone* compares ResNet depths to verify gains stem from the hyperbolic head rather than feature extraction. The full model (*d* = 256, *c* = 1.0, ResNet-18) achieves the best accuracy, confirming that hyperbolic geometry and the developmental prior contribute complementary benefits—the former provides a distance metric suited to hierarchical data, while the latter anchors prototypes to a biologically plausible manifold.

## IV. Results and Discussion

We evaluate HyperDev on 7-way few-shot developmental stage classification in *C. elegans*, demonstrating that hyperbolic geometry enables biologically meaningful embeddings while maintaining competitive performance despite severe class imbalance.

### A. Classifier Performance and Explainability

Table III presents accuracy across nine few-shot configurations (1/2/5-shot with 1/5/15 queries) over 500 episodes. HyperDev achieves competitive accuracy (76.9–88.3%) compared to ProtoNet (80.9–89.2%) and MatchingNet (81.2– 89.6%), with the 5-shot, 15-query configuration reaching 88.3%. RelationNet fails dramatically (14.3%), indicating metric-based approaches are more robust under extreme class imbalance. The 2–4% accuracy gap between HyperDev and top baselines is acceptable given the severe imbalance (L2: 6 samples, Dauer: 8 samples), representing a reasonable tradeoff for dramatically superior interpretability. For minority classes, per-class accuracy is inherently sensitive to individual predictions given the limited test instances available. This sensitivity is a property of the benchmark shared equally across all methods, rather than a weakness specific to HyperDev. Consequently, overall episodic accuracy across 500 episodes (Table III, Panel A) is the more statistically reliable metric at this data scale

#### Biological alignment

Table IV quantifies geometric– biological correspondence through multiple metrics. HyperDev exhibits significantly larger hyperbolic distances (Adjacent: 1.730, Non-Adjacent: 2.249) compared to Euclidean baselines (0.04–1.22), reflecting the Poincaré ball’s natural expansion. Critically, HyperDev’s mean radius of 0.583 (±0.100) differs significantly from baselines at 1.000 (*p <* 0.001, Welch’s *t*-test), indicating stages occupy the ball’s interior rather than the boundary. This radial distribution encodes developmental commitment: early stages with high plasticity position closer to the origin, while more developmentally committed or specialized stages are embedded toward the boundary.

**TABLE IV:** Quantitative analysis in terms of the explanability of models showing the performance of HyperDev compared to the baseline method.

| Method | Adjacent Distance | Non-Adjacent Distance | Separation Ratio | Mean Radius ± Std | Within-Stage Variance | Pearson $r$ ($p$) | Spearman $\rho$ ($p$) | Mantel $r$ ($p$) |
| --- | --- | --- | --- | --- | --- | --- | --- | --- |
| ProtoNet | 0.922 | 1.215 | 1.32 | 1.000 (0.000) | 0.021 | 0.187 (0.416) | 0.180 (0.434) | 0.187 (0.451) |
| MatchingNet | 0.989 | 1.217 | 1.23 | 1.000 (0.000) | 0.021 | 0.235 (0.306) | 0.281 (0.218) | 0.235 (0.398) |
| RelationNet | 0.041 | 0.064 | 1.58 | 1.000 (0.000) | 0.001 | 0.464 (0.034) | 0.471 (0.031) | 0.464 (0.046) |
| **HyperDev** | **1.730\*** | **2.249\*** | **1.30** | **0.583 (0.100)\*** | **0.015** | **0.669 (0.001)** | **0.570 (0.007)** | **0.669 (<0.001)** |

HyperDev exhibits consistently stronger geometric–biological correspondence than all baselines across correlation metrics. Pearson and Mantel coefficients are numerically identical due to linear correlation on vectorized distance matrices, while the Mantel test provides robust significance via permutation testing. Significant improvement over all baselines ( $p < 0.001$ , two-tailed Welch’s $t$ -test).

Fig 3 visualizes the correlation between biological distance (stage difference) and embedding distance. HyperDev achieves the strongest correlation (Pearson *r*=0.669, *p<*0.001; Spearman *ρ*=0.570, *p*=0.007), substantially outperforming ProtoNet (*r*=0.187, *p*=0.416), MatchingNet (*r*=0.235, *p*=0.306), and RelationNet (*r*=0.464, *p*=0.034). The Mantel test confirms this result (*r*=0.669, *p<*0.001) using non-parametric permutation testing. As shown in Fig 3(d), HyperDev exhibits a clear upward trend indicating that biologically distant stages are also far apart in embedding space, while Fig 3a–3c show baselines produce scattered points with weak or no correlation. Although Dauer is entered from L2 biologically, it represents a highly specialized alternative developmental fate; accordingly, it is plausible for Dauer to appear at a larger radius despite not being strictly ‘later’ in linear time. This strong correlation demonstrates that HyperDev embeddings preserve the natural biological ordering of developmental stages. When two stages are biologically distant (e.g., Egg vs. Adult), they are also far apart in embedding space; conversely, adjacent stages (e.g., L2 vs. L3) remain close. Euclidean baselines fail to capture this relationship effectively, with ProtoNet and MatchingNet showing nearly random correlation (*p >* 0.3).

**Fig. 3:**
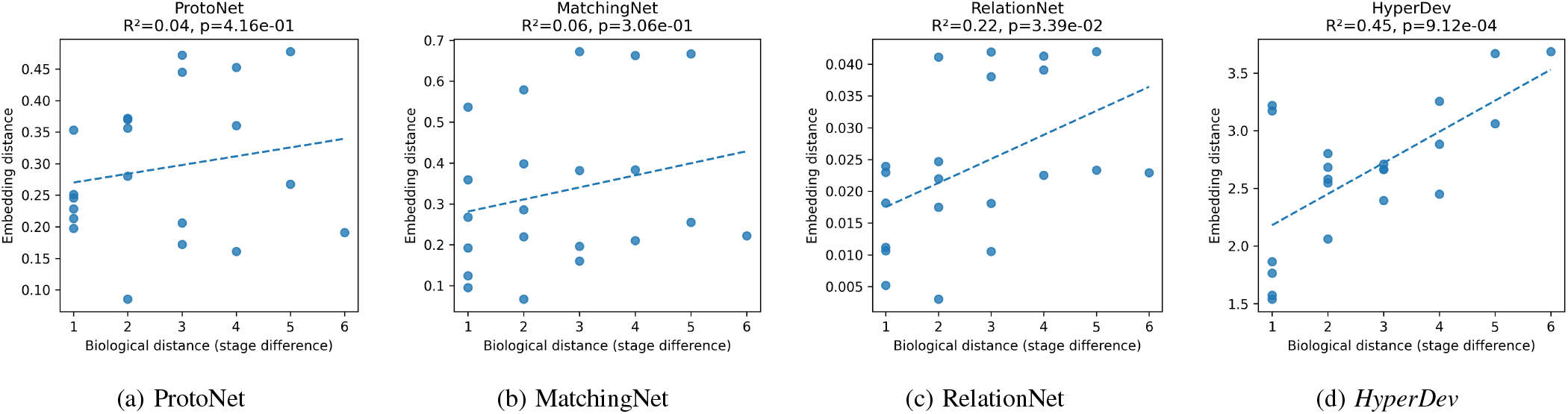
Biological vs. embedding distance correlation. HyperDev achieves significantly stronger alignment (*r* = 0.669, *p <* 0.001) than baselines, demonstrating superior capture of developmental hierarchy.

#### Geometric interpretability

Within-stage variance (0.015) demonstrates tight clustering comparable to baselines (0.001– 0.021), maintaining intra-class cohesion while achieving superior inter-class organization. The radial structure reveals meaningful hierarchy: developmental progression can be interpreted as a trajectory from the origin (early, plastic stages) toward the boundary (more committed or specialized outcomes), emerging naturally from hyperbolic geometry’s ability to embed tree-like structures. Error analysis shows majority of mistakes occur between adjacent stages (e.g., L2 ↔ L3), demonstrating biological plausibility—the model rarely confuses distant stages (e.g., Egg vs. Adult), respecting coarse-grained structure even when fine-grained classification is challenging.

### B. Discussion

#### Accuracy vs. interpretability trade-off

The compromise in 2–4% of HyperDev accuracy than the top baselines models is mainly due to the following three main factors:

- Flexibility is reduced by geometric constraints.
- The biological structures are prioritized by development over optimal decision boundaries.
- Severe class imbalance (L2: 6 samples, Dauer: 8)

This modest trade-off between accuracy and explainability results in a stronger biological correlation (*r*=0.669 vs. 0.187–0.464), with the majority of errors occurring between adjacent stages rather than distant ones, indicating biologically plausible failures.

#### Why hyperbolic geometry succeeds

The Poincaré ball presents two dimensions that display developmental commitment: position from the origin and stage characteristics through its angular position. Biological development begins with pluripotent cells that progress along distinct developmental trajectories. The system developed its structure without explicit supervision, producing meaningful representations that revealed biological relationships through their geometric structure.

#### Value for biological research

The model achieving 88% accuracy with explainability is more valuable than a 89% black-box model as it represents biologically grounded representations. HyperDev enables essential features, including visualization of developmental relationships and understanding of model decisions, that standard approaches lack, which could reveal unknown relationships in less-studied organisms.

#### Limitations and future work

Hyperbolic-awareness or learned developmental priors may close the accuracy gap while preserving explainability, but are limited to *C. elegans*, so extensions to other hierarchical biological problems (cell differentiation, phylogenetics) represent promising directions. Expanding the L2 and Dauer test sets beyond 6 and 8 instances respectively would reduce per-class accuracy sensitivity and enable more reliable confidence interval estimation for minority stages — a priority for future dataset curation.

## V. Conclusion

We introduced HyperDev, a hyperbolic few-shot learning framework for *C. elegans* developmental stage classification. Utilizing the Poincaré ball’s natural hierarchical structure, HyperDev achieves competitive accuracy (76.9–88.3%) with substantially superior biological interpretability (*r*=0.669 vs. 0.187–0.464). The learned embeddings reveal meaningful organization-with the majority of errors between adjacent stages. Our work establishes hyperbolic geometry as a principled framework for explainablebiological few-shot learning, where understanding representations is essential for discovery. Future work will explore hyperbolic-aware augmentation and extensions to hierarchical biological problems.

## Ethics Statement

The invertebrate nematode *Caenorhabditis elegans* was used for the experiments described in this study and does not require approval by the Institutional Animal Care and Use Committee (IACUC).

## Acknowledgements

The authors received partial support from the Missouri S&T Mary K. Finley Endowment, the Kummer Institute for Artificial Intelligence and Autonomous Systems, and Missouri S&T’s IGI funding. This work was also supported by NSF Award No. 2334884 under the project title “BRC-BIO: The Impact of Pheromone Signaling on *C. elegans* Population Dynamics,” and NSF Award No. 2420248 under the project title “EAGER: Solving Representation Learning and Catastrophic Forgetting with Adaptive Resonance Theory.” The views and conclusions contained in this document are those of the authors and should not be interpreted as representing the official policies, either expressed or implied, of the National Science Foundation or the U.S. Government. The U.S. Government is authorized to reproduce and distribute reprints for Government purposes notwithstanding any copyright notation herein.

## References

[1] Z. F. Altun and D. H. Hall, “Introduction,” WormAtlas, 2009.

[2] Z. Kocsisova, B. M. Egan, A. Scharf, X. Anderson, F. Pohl, A. Anderson, and K. Kornfeld, “How to measure, analyze, and interpret age-related changes in caenorhabditis elegans: lessons for mechanistic and evolutionary theories of aging,” Mechanisms of Ageing and Development, p. 112146, 2025.

[3] K. A. Caldwell, C. W. Willicott, and G. A. Caldwell, “Modeling neurodegeneration in caenorhabditis elegans,” Disease Models & Mechanisms, vol. 13, no. 10, p. dmm046110, 2020.

[4] L. P. O’Reilly, C. J. Luke, D. H. Perlmutter, G. A. Silverman, and S. C. Pak, “C. elegans in high-throughput drug discovery,” Advanced drug delivery reviews, vol. 69, pp. 247–253, 2014.

[5] J. E. Sulston, E. Schierenberg, J. G. White, and J. N. Thomson, “The embryonic cell lineage of the nematode caenorhabditis elegans,” Developmental biology, vol. 100, no. 1, pp. 64–119, 1983.

[6] T. Kaletta and M. O. Hengartner, “Finding function in novel targets: C. elegans as a model organism,” Nature reviews Drug discovery, vol. 5, no. 5, pp. 387–399, 2006.

[7] H. R. Horvitz, “Genetic control of programmed cell death in the nematode caenorhabditis elegans,” Cancer Research, vol. 59, no. 7 Supplement, pp. 1701s–1706s, 1999.

[8] A. Fire, S. Xu, M. K. Montgomery, S. A. Kostas, S. E. Driver, and C. C. Mello, “Potent and specific genetic interference by double-stranded rna in caenorhabditis elegans,” Nature, vol. 391, no. 6669, pp. 806–811, 1998.

[9] O. Vinyals, C. Blundell, T. Lillicrap, D. Wierstra et al., “Matching networks for one shot learning,” Advances in neural information processing systems, vol. 29, 2016.

[10] O. Ganea, G. Bécigneul, and T. Hofmann, “Hyperbolic neural networks,” in Advances in Neural Information Processing Systems, 2018, pp. 5345–5355.

[11] M. Nickel and D. Kiela, “Poincaré embeddings for learning hierarchical representations,” in Advances in Neural Information Processing Systems, 2017, pp. 6338–6347.

[12] Y. Liu, Z. He, and K. Han, “Hyperbolic category discovery,” in Proceedings of the Computer Vision and Pattern Recognition Conference (CVPR), June 2025, pp. 9891–9900.

[13] K. He, X. Zhang, S. Ren, and J. Sun, “Deep residual learning for image recognition,” in Proceedings of the IEEE Conference on Computer Vision and Pattern Recognition, 2016, pp. 770–778.

[14] A. Vaswani, N. Shazeer, N. Parmar, J. Uszkoreit, L. Jones, A. N. Gomez, Ł. Kaiser, and I. Polosukhin, “Attention is all you need,” in Advances in Neural Information Processing Systems, 2017, pp. 5998–6008.

[15] F. Zheng, C. Chen, and M. Aschner, “Neurotoxicity evaluation of nanomaterials using c. elegans: Survival, locomotion behaviors, and oxidative stress,” Current protocols, vol. 2, no. 7, p. e496, 2022.

[16] L. Byerly, R. Cassada, and R. Russell, “The life cycle of the nematode caenorhabditis elegans: I. wild-type growth and reproduction,” Developmental biology, vol. 51, no. 1, pp. 23–33, 1976.

[17] X. Karp, “Working with dauer larvae,” WormBook, vol. 2018, p. 1, 2018.

[18] S. Bhattiprolu, “Digitalsreeni image annotator,” https://github.com/bnsreenu/digitalsreeni-image-annotator, 2024, accessed: 2025-12-01.

[19] R. Lints and D. H. Hall, “Wormatlas hermaphrodite handbook-reproductive system-egg-laying apparatus,” WormAtlas, 2004.

[20] J. E. Sulston and H. R. Horvitz, “Post-embryonic cell lineages of the nematode, caenorhabditis elegans,” Developmental biology, vol. 56, no. 1, pp. 110–156, 1977.

[21] J. Snell, K. Swersky, and R. Zemel, “Prototypical networks for few-shot learning,” Advances in neural information processing systems, vol. 30, 2017.

[22] F. Sung, Y. Yang, L. Zhang, T. Xiang, P. H. Torr, and T. M. Hospedales, “Learning to compare: Relation network for few-shot learning,” in Proceedings of the IEEE conference on computer vision and pattern recognition, 2018, pp. 1199–1208.

